# *Marchantia polymorpha* as a simple platform for plant-based production of functional nanobodies

**DOI:** 10.64898/2026.06.22.733889

**Authors:** Sze Wai Tse, Facundo Romani, Fernando Guzman-Chavez, Eftychios Frangedakis, Jim Haseloff

## Abstract

Recombinant proteins have transformative potential in biomedicine, but their production is often costly and carries contamination risks. Plants offer an attractive alternative, with low growth costs and reduced pathogen risk, yet their slow growth cycles limit their use for rapid protein engineering. Here, we establish *Marchantia polymorpha*, a genetically tractable liverwort with a short life cycle, as a new platform for recombinant protein production. Using stable Agrobacterium-mediated transformation, we expressed an anti-mCherry nanobody fused to the fluorescent protein mTurquoise2 with different purification tags. Expression levels reached up to ∼120 µg/g fresh weight, and nanobody functionality was validated through a microscopy-based bead-binding assay. This yield rivals that of established systems such as *Nicotiana benthamiana*. Our results position *M. polymorpha* as a scalable, safe, and efficient chassis for protein engineering, with broad potential for applications in synthetic biology. This work opens the door to exploiting liverwort biotechnology for fast, cost-effective, and biosafe production of valuable recombinant proteins.

**Graphical Table of Contents:** 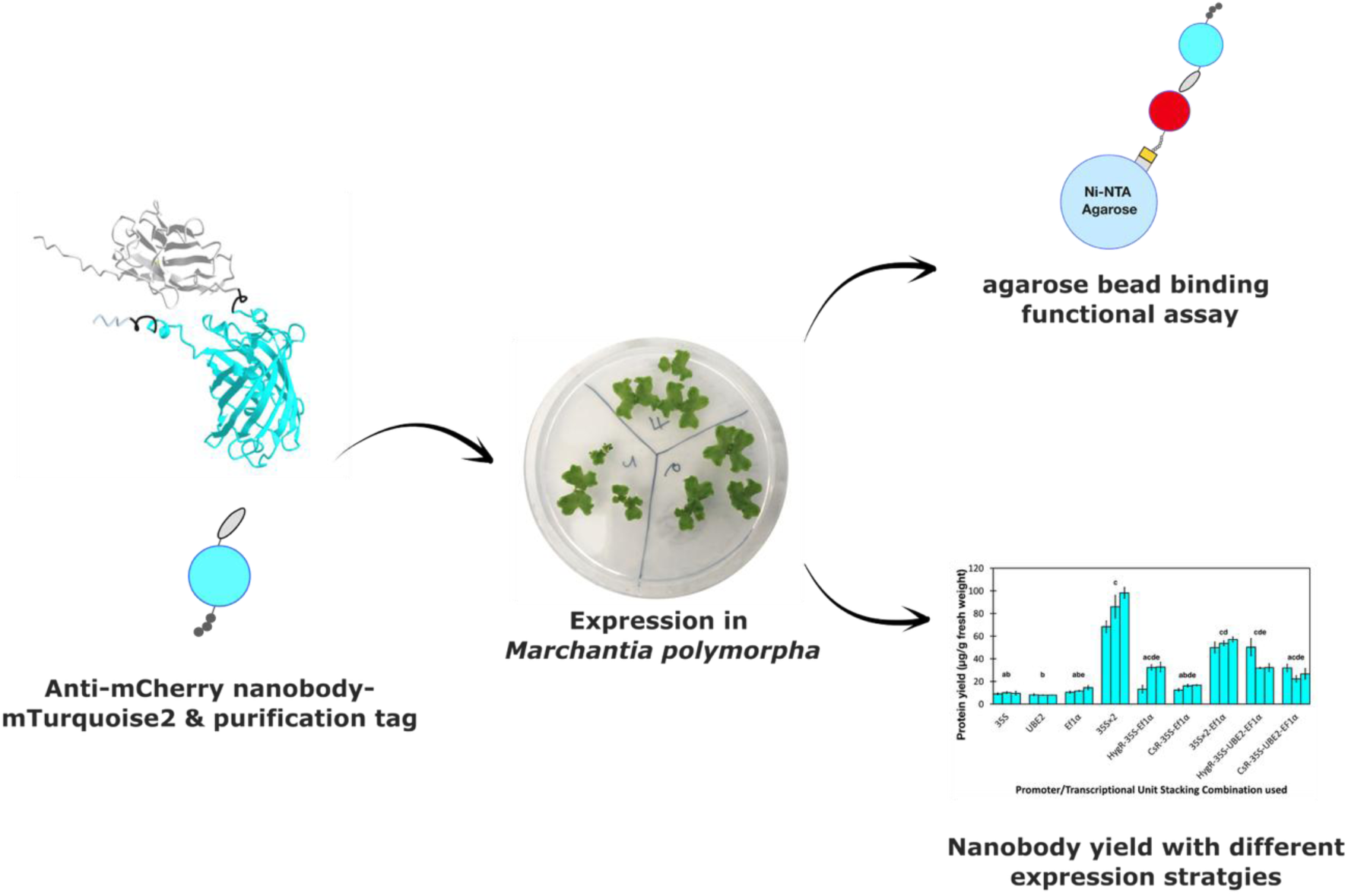

## INTRODUCTION

Production of recombinant protein, often expressed from recombined or engineered DNA in a heterologous host^1^, is important for making biologics for therapeutic and diagnostic purposes^2,3^. Currently, the majority of applications are based on bacterial systems such as *Escherichia coli* (*E. coli*)^4^ and mammalian systems such as the Chinese hamster ovary (CHO) cells ^5^. Plants present strong potential for recombinant protein production due to their relative ease of growth, potential for scale-up, and low risk of contamination with human pathogens^6^ comparing to bacterial and mammalian systems. compared to bacterial and mammalian systems. However, stable expression of recombinant protein is time consuming in plants. Even transient expression in popular systems such as *Nicotiana benthamiana*^7^ can take up to 2-4 weeks^8^ and is labour- and capital-intensive to perform at large scale, partially explaining the lack of extensive adoption of plant-based recombinant proteins.

*Marchantia polymorpha*, an emerging plant model for synthetic biology^9^, has a series of traits that can help to overcome to these bottlenecks. It has a rapid life cycle, does not require sophisticated equipment for growth and stable transformation can be performed within a timescale of weeks, as opposed to months or longer in flowering plants. Growth of plant material either by vegetative propagation of thallus, or million-fold amplification by sexual production of spores is straightforward, presenting great potential for scale-up^10^. Recently a series of standardised genetic tools for the efficient construction of recombinant protein expression cassettes for both nuclear and chloroplast expression has become available, which further streamline the recombinant protein production process in *M. polymorpha*^9,11,12^. As a proof of concept of the suitability of *M. polymorpha* for the production of recombinant proteins, we demonstrated the potential to obtain over 400 µg/g of fresh weight mTurquoise2 protein after chloroplast transformation^13^.

However, the suitability of *M. polymorpha* for the large-scale production of a wider range of compounds with commercial value has not been tested yet. To explore this, we selected nanobodies, corresponding to the antigen binding domains of single-chain antibodies from camelids^14^, present an interesting test-case. They are ∼15kDa in size, 10% the size of convention monoclonal antibodies^15^, yet retain high binding specificity. They are routinely expressed in *E. coli*^16^ and have a market size of 368.6 million US dollars for applications in therapeutics and diagnostics, particularly after the COVID-19 pandemic^17^. Expression levels were comparable to those achieved when nanobodies have been expressed in plant systems, from *Nicotiana benthamiana*^18,19^ to Arabidopsis seeds^20,21^ and potato^22^. Again, these are flowering plant models with month- to year-long life and experimental cycles for time-efficient prototyping of nanobody production, therefore presenting the need for a faster growing system such as Marchantia to facilitate development of better expression strategies for nanobodies.

In this study, we tested the speed and efficiency of the *M. polymorpha* system and demonstrated successful expression and functional verification of an anti-mCherry nanobody-Turquoise2 fusion protein (LaM4-mTurquoise2) using stable nuclear transformation in *M. polymorpha*. The impact of purification tags on yield and purification efficiency was also tested.

## RESULTS

### Nanobody expression candidate and microscopy-based assay

We selected the anti-mCherry nanobody LaM4 with its published amino acid sequence^23^ and structure^24^ as the candidate for expression in *M. polymorpha*. This nanobody has been shown to bind to its target when tagged with horseradish peroxidase (HRP) for fluorescence signal amplification and a polyhistidine (7xHis) tag for protein purification^25^. We fused it to the *Aequoria victoria* GFP-derived cyan fluorescent protein mTurquoise2 that gives bright signal in *M. polymorpha*^9^ and has a sequence distinct from the dsRed-derived mCherry^26^ to negate binding to LaM4. A C-terminal 7x-histidine tag was also included to facilitate purification and visualisation of binding between the nanobody and its target (Figure 1b).

**Figure 1:**
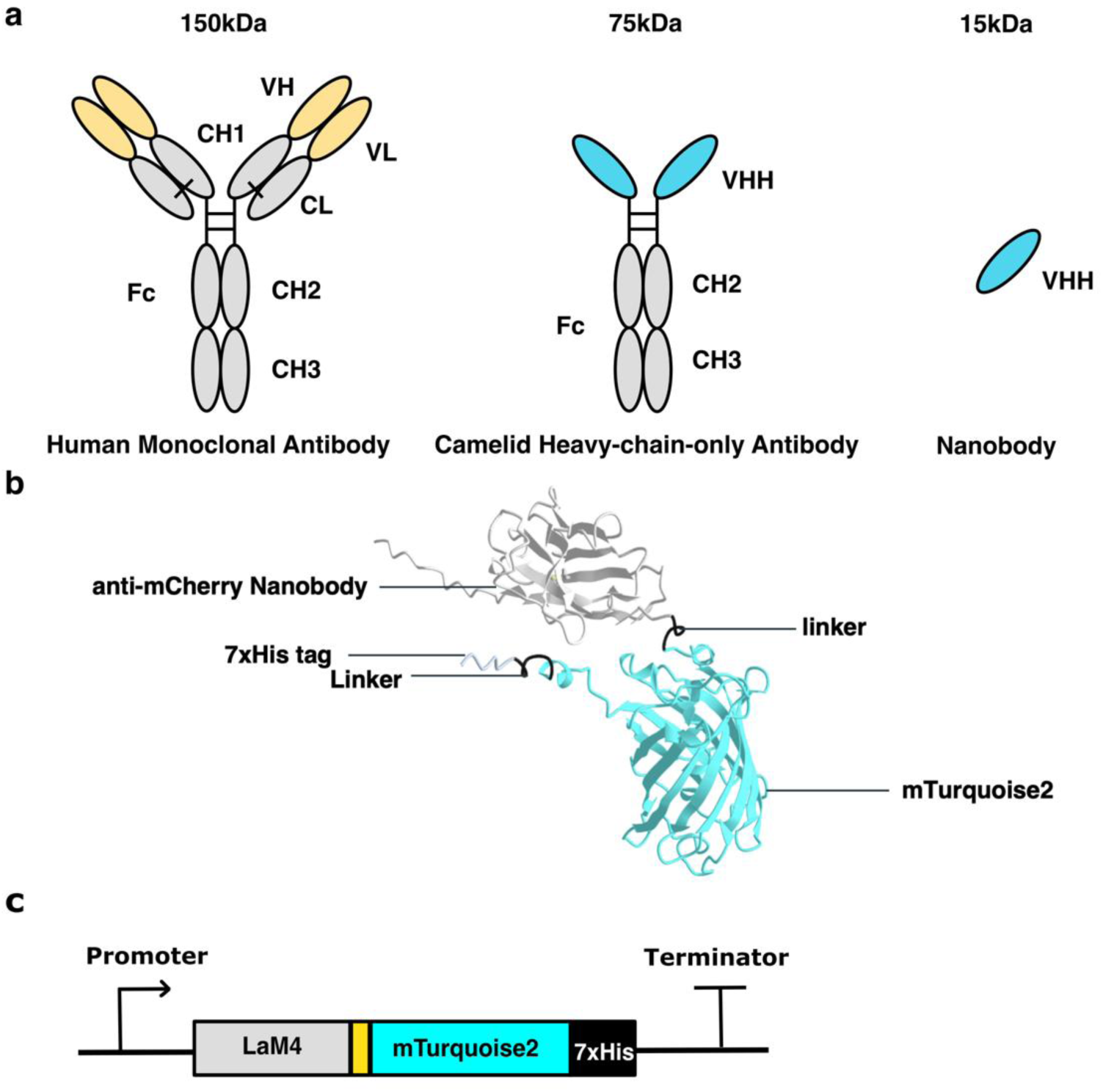
Design of the anti-mCherry-nanobody-mTurquoise2 fusion protein for expression in M. polymorpha. **a**: A schematic description of nanobody in relation to a human monoclonal antibody with both heavy and light chains and a Camelid-heavy chain only nanobody. Fc: Fragment crystallizable. CH1,2,3: constant domain 1,2,3 of the heavy chain. CL: constant domain of the light chain. VH: variable domain of the heavy chain. VL: variable domain of the light chain. VHH: Variable domain of heavy-chain-only antibodies, also called nanobodies. Figure adapted from^15,27^ **b**: A 3D structure generated with AlphaFold2^28^ for the anti-mCherry nanobody (LaM4) fused to mTurquoise2 and the 7x-Histidine tag (LaM4-Turquoise2-7xHis). **c:** Construct map for LaM4-mTurquoise2-7xHis.

To test for correct folding and functionality of LaM4-mTurquoise2-7xHis fusion, we first expressed and purified the protein in *E. coli*. When the purified LaM4-mTurquoise2-7xHis was loaded onto nickel-nitrilotriacetic acid (Ni-NTA) agarose beads and incubated with its antigen mCherry, fluorescence signals from both mCherry and mTurquoise2 were visible on the beads under a confocal microscope (Figure 2d), in contrast to the controls (Figure 2a-c). LaM4-Turquoise2-7xHis was captured through interaction between Ni-NTA on the agarose beads and the 7x-histidine tag, while the mCherry binding required attachment through interaction with LaM4-Turquoise2-7xHis, demonstrating functional binding by the latter.

**Figure 2:**
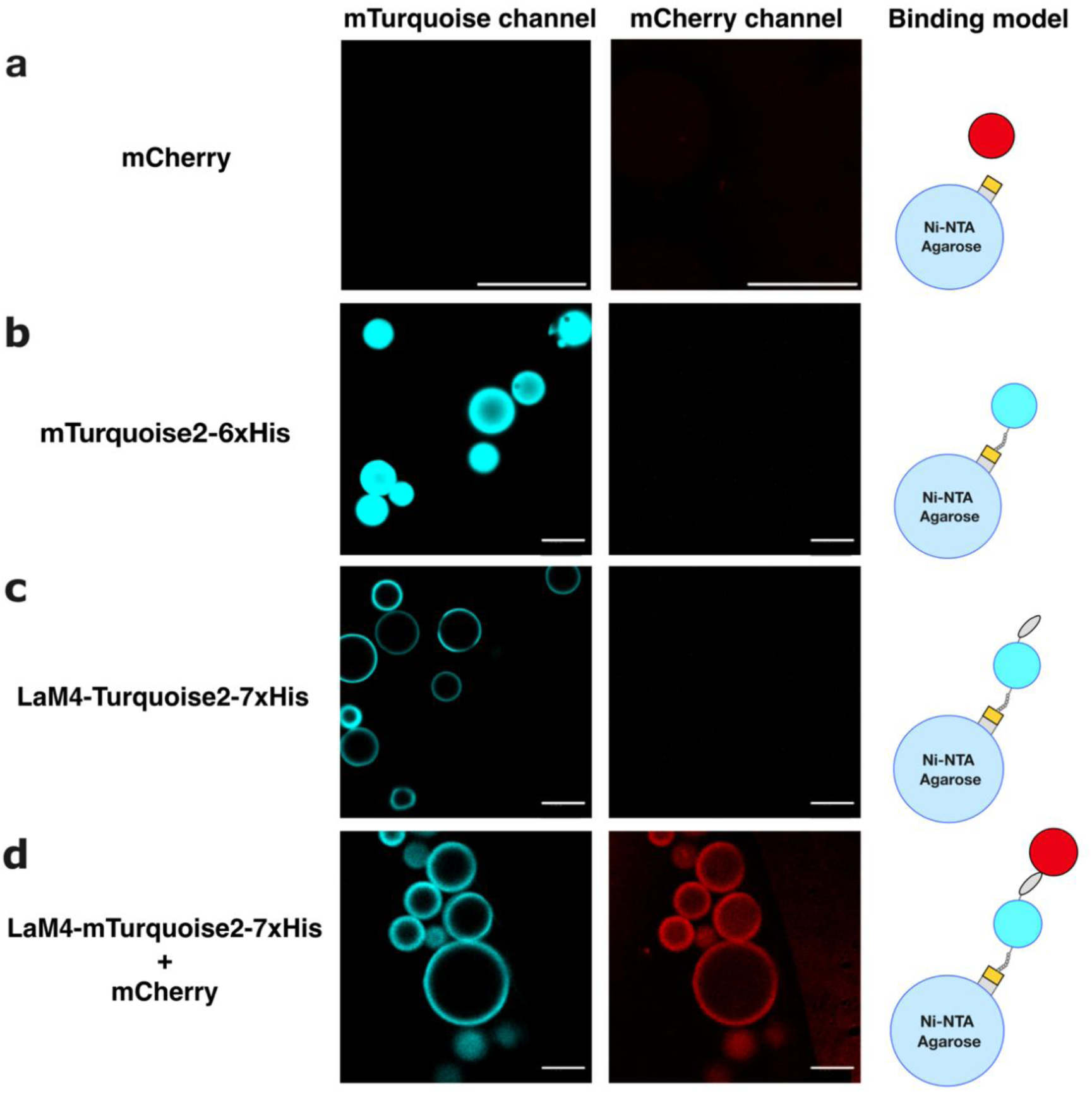
Ni-NTA agarose bead binding assay used to verify the binding between LaM4-Turquoise2-7xHis produced in E. coli and its antigen, mCherry. Each row shows the confocal microscopy images of protein(s) incubated with Ni-NTA agarose beads, alongside their corresponding binding models. **a**: tag-less mCherry. **b**: mTurquoise2-7x-His. **c**: LaM4-mTurquoise2-7xHis. **d**: LaM4-mTurquoise2-7xHis and mCherry. mTurquoise2 signal shown as cyan and mCherry signal shown in red. Scale = 100µm.

### Expression of LaM4-mTurquoise2-7xHis in *M. polymorpha*

To find the best gene design for maximal recombinant protein yield in *M. polymorpha*, we expressed the nanobody under various constitutive promoters available from the OpenPlant kit^9^ as single transcription units^12^ and stacked multiple nanobody expression transcription units in one construct^29^ (Figure 3a), then estimated the yield of LaM4-mTurquoise2-7xHis from each line by accumulated mTurquoise2 fluorescence^12,13^. Amongst all single, dual and triple transcriptional-unit(s) lines, pro35S×2: LaM4-mTurquoise2-7xHis gave the markedly highest yield at ∼100 µg/g fresh weight. Interestingly, this yield was also higher than those obtained from dual- and triple-transcriptional-units expressing plants, even where the strong pro35S×2 and proMp*EF1*α promoters were used to drive expression of LaM4-mTurquoise2-7xHis. The yields between hygromycin or chlorsulfuron resistant transformants with dual and triple stacked transcription units were not significantly different from each other, indicating that the selection marker does not lead to a significant difference between plants (Figure 3b). Accumulated biomass from all lines were significantly less than that in WT, except pro35S: LaM4-mTurquoise2-7xHis, proMp*EF1*α: LaM4-mTurquoise2-7xHis and pro35S×2: LaM4-mTurquoise2- proMp*EF1*α: LaM4-mTurquoise2-7xHis dual expression cassette lines (Figure 3c, d). This indicates as consistent with our previous publication, higher level of accumulation of recombinant protein levels is correlated with higher level of growth penalty^12^.

**Figure 3:**
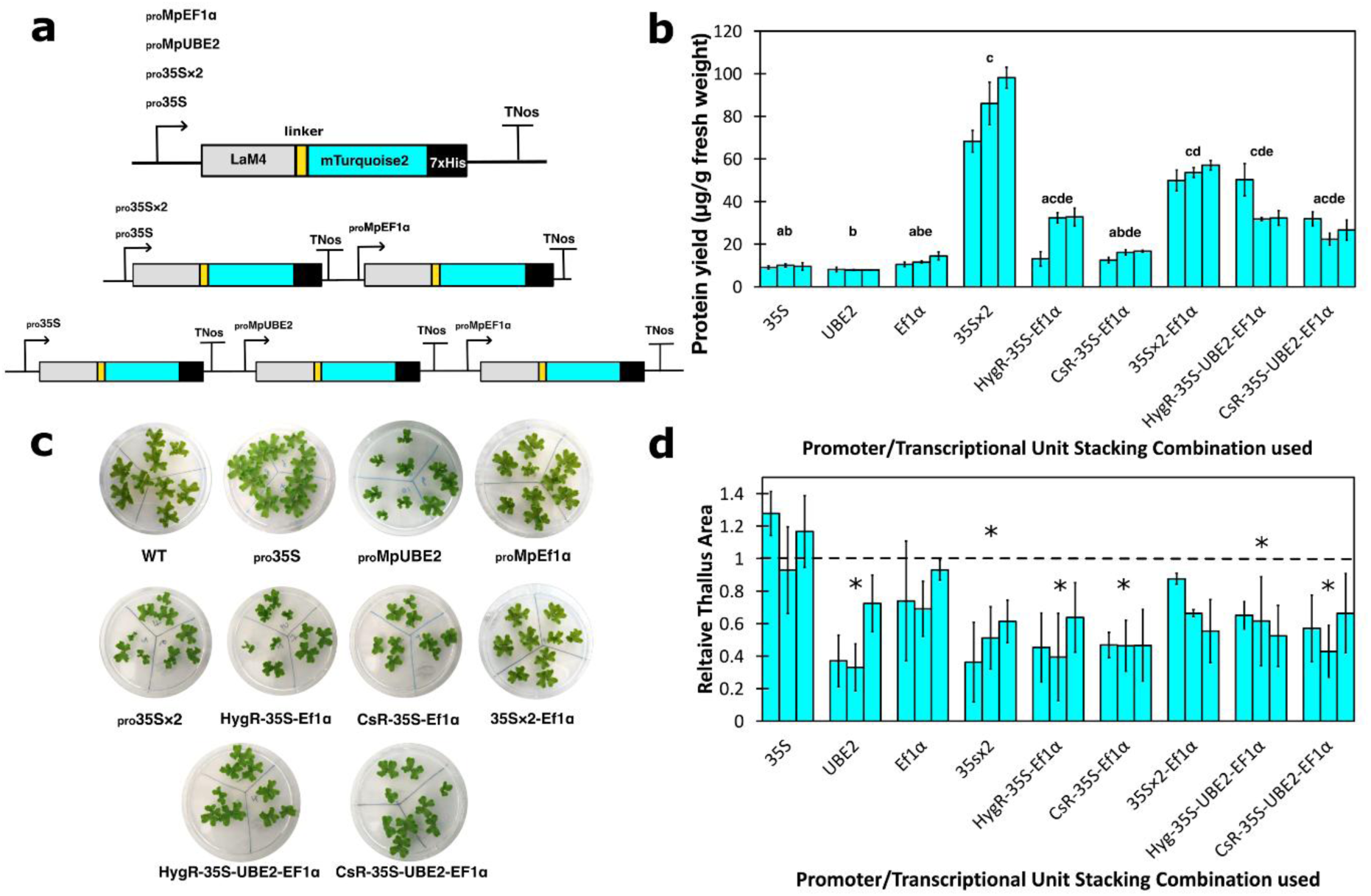
Expression level of LaM4-mTurquoise2-7xHis and relative size estimation of plants from different in different promoter:LaM4-mTurquoise2-7xHis lines. **a**: Schematic representation of different promoter:LaM4-mTurquoise2-7xHis constructs. **b**: LaM4-mTurquoise2-7xHis expressed in different promoter:LaM4-mTurquoise2-7xHis in 3-week-old plants, in µg/g fresh weight. **c**: Images of 2-week-old WT and different promoter:LaM4-mTurquoise2-7xHis plants. **d**: Relative thallus size (the ratio of sizes of promoter:LaM4-mTurquoise2-7xHis plants: average size of WT plants; dash line indicates WT size) of different promoter line plants (2-week-old). In **b** and **d**, each bar represents an independent transformant for that promoter construct, and error bars represents standard deviation between 3 biological replicates. Letters/asterisk above the bars indicate statistically significant differences between the different promoter:LaM4-mTurquoise2-7xHis (Dunn’s Test; p < 0.05). HygR and CsR prefixes attached to the names of the constructs indicate the plant selection markers used, for comparison of hygromycin and chlorsulfuron resistance. If no selection marker has been noted, hygromycin resistance selection marker has been used.

We also co-transformed multiple stacked transcription units into *M. polymorpha*, as this approach could improve recombinant protein yield^29^. The expression levels of LaM4-mTurquoise2-7xHis in plants co-transformed with 2 and/or 3 stacked transcription units (Figure S1a) are not statistically significantly different from those with a single transcription unit (Figure 3, S1b), all approximately half the yield obtained in pro35S×2: LaM4-mTurquoise2-7xHis. The sizes of the co-transformed plant lines were statistically significantly smaller than WT plants, except for plants co-transformed with triple expression cassette plasmids. (Figure S1c, d).

To test for the effect of different tags on the yield of recombinant protein in *M. polymorpha* as well as test for easier purification in *M. polymorpha*, we chose two alternative tags to test for purification of LaM4-mTurquoise2 from *M. polymorpha*: the FLAG short peptide tag that binds to the M2 antibody^30^ and the Car9 silica-binding short peptide tag^31^. FLAG tag in a tandem of three (3xFLAG) had been successfully used previously for purification of protein produced in *M. polymorpha*^32^, while the Car9 tag allows for purification using cheaper resins. We built two constructs for Car9, one with (cCar9) and without (Car9) a domain cleavable by the *E. coli* membrane enzyme OmpT for tag removal (Figure 4a). 3X-FLAG tagged LaM4-mTurquoise2 gave the highest yield amongst all transgenic plants expressing alternative tags (Figure 4b) and produced no statistically significant negative effect on plant growth as opposed to other lines (Figure 4c, d).

**Figure 4:**
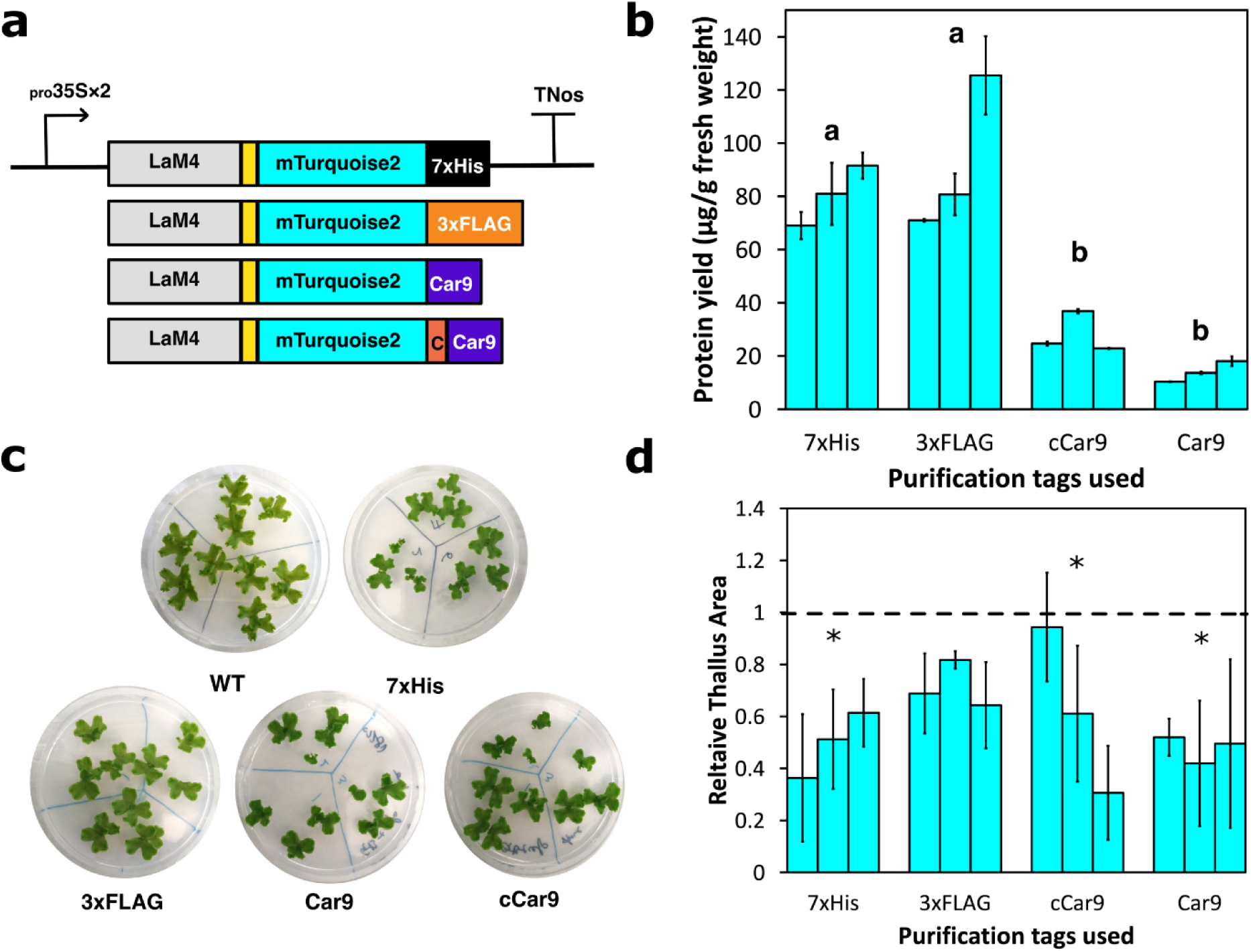
Expression level of LaM4-mTurquoise2 with different tags and relative size estimation of plants in different LaM4-mTurquoise2 alternatively tagged lines. **a**: Schematic representation of different LaM4-mTurquoise2 alternative tag constructs. **b**: LaM4-mTurquoise2 expressed in different LaM4-mTurquoise2 alternative tag in 3-week-old plants, in µg/g fresh weight. **c**: Images of 2-week-old WT and LaM4-mTurquoise2 alternative tag plants. The plates were divided into three portions, and on each portion grew 3 gemmae derived from the same independent transformant of that promoter construct. **d**: Relative thallus size (the ratio of sizes of LaM4-mTurquoise2 alternative tag plants: average size of WT plants; dash line indicates WT size) of different LaM4-mTurquoise2 alternative tag plants (2-week-old). In **b** and **d**, each bar represents an independent transformant for each construct, and error bars represents standard deviation between 3 biological replicates. Asterisks above the bars indicate statistically significant differences between the different LaM4-mTurquoise2 alternative tag constructs and WT plants (Dunn’s Test; p < 0.05).

The 3xFLAG-tagged LaM4-mTurquoise2 could be captured only on anti-FLAG agarose beads (Figure 5b), but not on anti-His Ni-NTA agarose beads (Figure 5c) and the anti-FLAG beads binds specifically to FLAG-tagged LaM4-mTurquoise2 only (Figure 5d). While FLAG-tagged LaM4-mTurquoise2 binding to mCherry could not be visualised on anti-FLAG agarose beads, similar to the *E. coli* assay showed in Figure 2, in a different setting 3xFLAG-tagged LaM4-mTurquoise2 can be immobilised to mCherry-7xHis bound to Ni-NTA beads (Figure 5a), which the nanobody on its own did not interact with (Figure 5c), demonstrating the functionality of the plant-produced 3xFLAG-tagged LaM4-mTurquoise2. No mTurquoise2 signal was visible when mCherry-7xHis was incubated with WT plant extract and Ni-NTA beads, or with mTurquoise2 (Figure S2), demonstrating that the signal observed in experimental samples was induced over background.

**Figure 5:**
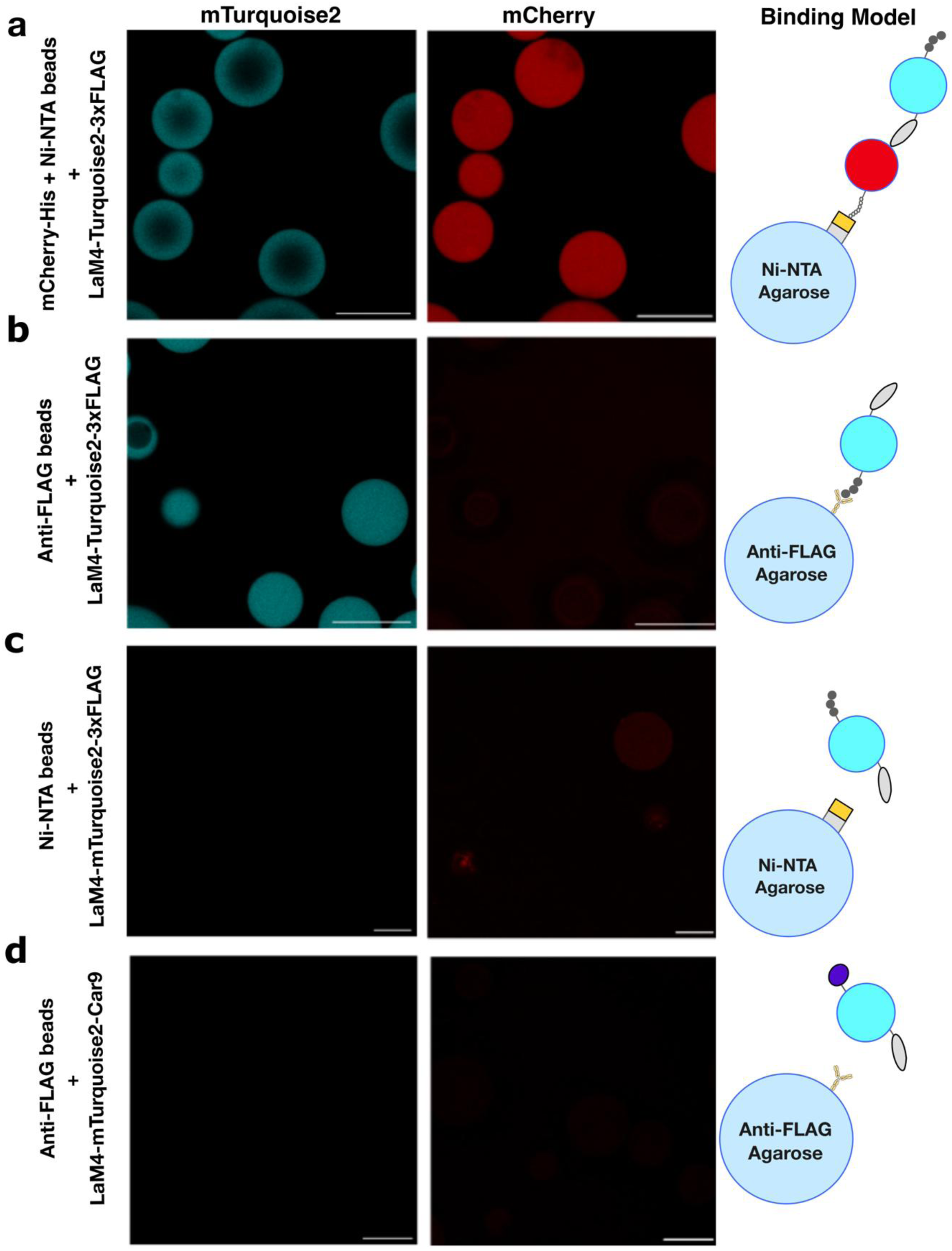
Bead binding assay with LaM4-mTurquoise2-3xFLAG binding to its intended target, mCherry, and anti-FLAG agarose beads, alongside respective negative controls. **a:** LaM4-mTurquoise2-3xFLAG incubated with mCherry-7xHis and Ni-NTA agarose beads. **b:** LaM4-mTurquoise2-3xFLAG incubated with anti-FLAG agarose beads. **c**: LaM4-mTurquoise2-3xFLAG incubated with Ni-NTA agarose beads. **d:** LaM4-mTurquoise2-Car9 incubated with anti-FLAG beads as negative control. mTurquoise2 signal shown as cyan and mCherry signal shown in red. Scale = 100µm.

## DISCUSSION

To date, different plant species have been shown to produce recombinant proteins, such as antibodies and nanobodies. Most if not all the systems used, such as *N. benthamiana*, *A. thaliana* and lettuce, are flowering plants with growth and experimental cycles which span months and even years, making testing and scaling up of recombinant protein production challenging. *M. polymorpha* is a fast-growing model liverwort with established genetic engineering toolkit and offers great potential for accelerating the engineering and production of recombinant proteins including nanobodies. In this paper, we have designed a LaM4-mTurquoise2-7xHis fusion protein as a prototype for exploring *M. polymorpha*’s potential as a testbed, together with a quick microscopy-based assay to confirm protein functionality.

The highest yield of LaM4-mTurquoise2 obtained was ∼120 µg/g fresh weight in LaM4-mTurquoise2-3xFLAG plants. This is comparable with the yields of single-domain antibody obtained in *Nicotiana tabacum* cv. Xanthi (136 μg/g of fresh leaf weight^33^) and nanobodies in *N. benthamiana* (129.50 mg/kg fresh weight^34^).

The yields in transgenic lines with stacked transcription units are lower than in the highest-expressing single-promoter lines, such as the pro35S×2: LaM4-mTurquoise2-7xHis and pro35S×2: LaM4-mTurquoise2-3xFLAG lines, different from what was observed in sugarcane^29^. This could be partially due to post-transcriptional gene silencing, known to lead to reduced recombinant protein yield in transgenic plants^35^. Potentially, this could be countered by co-transformation of the gene of interest with P19 repressor for gene silencing^36^, or by knocking out genes coding for proteins on the post-transcriptional gene silencing short-RNA metabolic pathway, such as RDR6^37^ and DCL2/4^38^.

We have previously demonstrated that high level of recombinant protein accumulation can be associated with decrease in plant biomass, but mTurquoise2 can accumulate in the cytosol to ~60 µg/g fresh weight in *M. polymorpha* with minimal detriment to plant growth^12^. This is consistent with our observation that plant growth is not significantly impacted in LaM4-mTurquoise2-3XFLAG tagged plants and suggests that *M. polymorpha* is a good system for reaching yields of recombinant proteins comparable to other plant systems.

Overall, our study demonstrates that *M. polymorpha* is a promising fast and efficient platform for recombinant protein production, achieving nanobody yields on par with established flowering plant systems. Its rapid growth, ease of genetic manipulation, and ability to accumulate high protein levels with minimal impact on biomass make it a promising alternative for scalable, plant-based biomanufacturing. While further optimization is needed for improving yield, costs, and downstream processing, the attributes of *M. polymorpha* position it as a potentially valuable testbed for molecular farming of recombinant proteins in plants.

## METHODOLOGY

### Plant Material and Growth Conditions

*Marchantia polymorpha subs. rudelaris* accessions Cam-1 (male) and Cam-2 (female) were grown as previously described^9^.

### Plasmid assembly

L1 and L2 backbones, as well as L0 promoter and terminator constructs were either from the OpenPlant Kit^9^ or from the previously published promoter collection^39^. L3-acceptor (pBy_10) was described before^12^. The coding sequence for LaM4 taken from Addgene plasmid #106410^25^, codon optimised using Geneious Prime for *E. coli* K-12 strain with default parameters and synthesised as gBlocks^TM^ (Integrated DNA Technologies) with BsaI cut sites and corresponding overhangs.

For the LaM4-mTurquoise2-7xHis construct, the mTurquiose2-7xHis fragment was amplified from the L0-mTurquoise2-CDS12 part^9^ using primers mTurquoise2-7xHis-F and mTurquoise2-7xHis-R. For the purification tag constructs, an mTurquoise2 fragment was PCR amplified from the L0-mTurquoise2-CDS12 part^9^ using primers mTurquoise2-7xHis-F and mTurquoise2-CDSI-R. Sequences coding for the purification tags were ordered as self-annealing primer pairs (Table S2). The gBlocks, PCR fragments and/or the primer dimer were cloned into L1 acceptors (pCk2, pCk3, pCk4) or directly into pBy_10, together with parts from the OpenPlant Kit^9^. For two- and three-transcriptional unit constructs, L1 vectors were assembled into L2 acceptor pCsA. Type-IIS cloning was performed as previously described^12^.

For bacterial expression, the pJL1-mCherry was a gift from Michael Jewett (Addgene plasmid #102629)^40^, and pCRB SREI6his was a gift from Christian Boehm^41^. The pmCherry-7xHis PCR fragment was amplified from pJL1-mCherry using primers mCherry-7xHis-F and mCherry-7xHis-R. The pmTurquoise2 PCR fragment was amplified from the L0-mTurquoise2-CDS12 part^9^ using primers mTurquoise2-F and mTurquoise2-R. PCR fragments were then cloned into pEPQD0KN0025-pJL1-atg-acceptor backbone^42^. The full list of plasmids used is available in Table S1, and all primers are listed in Table S2.

### E. coli protein expression and purification

Sequence-confirmed plasmids were transformed into BL21 Star^TM^ (DE3) pLysS One Shot^TM^ Chemically Competent *E. coli* (Thermo Fisher) according to the manufacturer’s instructions. Overnight cultures for each plasmid (50mL) were used to inoculate 250mL of LB medium supplemented with kanamycin and grown in 2.5mL glass flasks at 37°C and 220 rpm. Cultures were induced for T7 polymerase expression by addition of IPTG to a final concentration of 1mM, then grown for 5 hours at 37°C at 220 rpm. Cells were harvested by centrifugation at 6000g for 15 minutes at 4°C.

Harvested cells harbouring the non-tagged mCherry were disrupted using the lysis buffer from the Ni-NTA Fast Start Kit (Qiagen), with lysozyme and detergent supplied according to the manufacturer instructions. The lysates (crude extract of fluorescent proteins) were harvested by centrifugation at 14000g for 30 minutes at 4°C and stored as 1mL aliquots for further use.

For his-tagged LaM4-mTurquoise2-7xHis produced in *E. coli*, purification was performed according to manufacturer’s instructions for the Ni-NTA Fast Start Kit (Qiagen). Final elution fractions were collected and stored as 0.2mL aliquots.

### Bead binding assay

To test the functionality of the nanobodies with target proteins, we adapted a bead affinity detection protocol^43^ but relied solely on native fluorescence from the tags and targets^44^. We used 20µL of HisPur^TM^ Ni-NTA Superflow Agarose (Thermo Fisher) for binding to 50µL of nanobody extract and 50µL of fluorescent proteins (1-2mg/µL). All buffers were made according to the manufacturer’s instructions. The beads were incubated on an end-to-end rotator with the nanobodies and proteins premixed with 100µL of equilibration buffer at 4°C for 30 minutes and washed for 4 times using 100µL of wash buffer each time. The bead-nanobody-fluorescent protein slurries were stored in 100μL PBS (pH 7.4, Thermo Fisher).

For binding assays, plant extracts were made to 200µL with extraction buffer to give similar concentrations (∼150 µg/mL) of LaM4-mTurquoise2-7xHis/-3xFLAG/-Car9, with 20µL mCherry in *E. coli* crude extract and 20µL of HisPur^TM^ Ni-NTA Superflow Agarose (Thermo Fisher) or ANTI-FLAG® M2 Affinity Gel (A2220, Merck). Incubation time and wash-steps were performed the same as for bacterial samples.

For purification of anti-FLAG proteins, all assays were performed according to manufacturer’s protocol using ANTI-FLAG® M2 Affinity Gel (A2220, Merck) before the elution of LaM4-3xFLAG from the agarose beads. Bead-LaM4 slurries were subjected to confocal microscopy as described below.

### Laser Scanning Confocal Microscopy

20μL of the slurries from bead-binding assays were put into a 65μL Gene Frame (Thermo Fisher, AB0577) on a glass slide with a coverslip before imaging. Images were acquired on an upright Leica SP8X confocal microscope equipped with a 460–670 nm supercontinuum white light laser, 2 CW laser lines 405 nm, and 442 nm, and 5 channel spectral scanhead (4 hybrid detectors and 1 PMT). Imaging was conducted using either a 20× air objective (HC PL APO 20×/0.75 CS2) or a 40× water immersion objective (HC PL APO 40×/1.10 W CORR CS2). Excitation laser wavelength and captured emitted fluorescence wavelength window were as follows: for mTurquoise2 (442 nm, 460–485 nm) and for mCherry (587nm, 600-620nm); mCherry was imaged in separate sequential scans from mTurquoise2.

### Plant transformation, protein extraction and analysis

Plants were transformed using the Agrobacterium-mediated protocol for transformation in 6-well plates as described previously^39^. *M. polymorpha* protein extraction, yield and plant size estimation, and data handling was performed as previously described^12^.

## Supporting information

Additional information on experimental details, plasmids and primers used

## SUPPORTING INFORMATION

Additional information on experimental details, plasmids and primers used available as a PDF document.

## ACKNOWLEDGEMENT

We wish to thank Ignacy Bonter for helping troubleshooting protein extraction from *M. polymorpha* as well as confocal imaging, William Boxall for helping with statistical analyses using R, and Lei Hua for the gift of ANTI-FLAG® M2 Affinity Gel (A2220, Merck).

This work was funded as part of the BBSRC/EPSRC OpenPlant Synthetic Biology Research Centre Grant BB/L014130/1 to J.H., BBSRC BB/F011458/1 for confocal microscopy, BBSRC BB/T007117/1 to J.H. SWT was funded by the Doris Zimmern HKU-Cambridge Hughes Hall Scholarship. F.R. is a Leverhulme Early Career Fellow (ECF-2023-534) funded by the Leverhulme Trust and the Isaac Newton Trust (23.08(f)).

## AUTHOR CONTRIBUTIONS

JH helped SWT design the study and supervised the project. SWT and FGC did all *E. coli* protein expression, extraction, and purification. SWT did all plant experiment, with the help of FR and EF on transformation, plant protein extraction and troubleshooting in other aspects. All authors discussed the results and contributed to the final manuscript.

## DATA AVAILABILITY

All promoter and terminator L0 plasmids, and L1 and L2 backbones are from the OpenPlant kit^9^. L3 acceptor is as published before^12^. LaM4 Nanobody amino acid sequence is as previously published^25^. Primers are listed in the supplementary information.

## REFERENCES

(1) Overton, T. W. (2014) Recombinant protein production in bacterial hosts. Drug Discov. Today 19, 590–601.

(2) Miller, H. K., and Kersh, G. J. (2020) Analysis of recombinant proteins for Q fever diagnostics. Sci. Rep. 10, 20934.

(3) de Oliveira, A. L. G., Fraga, V. G., Sernizon-Guimarães, N., Cardoso, M. S., Viana, A. G., Bueno, L. L., Bartholomeu, D. C., da Silva Menezes, C. A., and Fujiwara, R. T. (2020) Diagnostic accuracy of tests using recombinant protein antigens of Mycobacterium leprae for leprosy: A systematic review. J. Infect. Public Health 13, 1078–1088.

(4) Rosano, G. L., Morales, E. S., and Ceccarelli, E. A. (2019) New tools for recombinant protein production in Escherichia coli: A 5-year update. Protein Sci. 28, 1412–1422.

(5) Tihanyi, B., and Nyitray, L. (2020) Recent advances in CHO cell line development for recombinant protein production. Drug Discov. Today Technol. 38, 25–34.

(6) Burnett, M. J. B., and Burnett, A. C. (2020) Therapeutic recombinant protein production in plants: Challenges and opportunities. PLANTS PEOPLE PLANET 2, 121–132.

(7) Bally, J., Jung, H., Mortimer, C., Naim, F., Philips, J. G., Hellens, R., Bombarely, A., Goodin, M. M., and Waterhouse, P. M. (2018) The Rise and Rise of *Nicotiana benthamiana* : A Plant for All Reasons. Annu. Rev. Phytopathol. 56, 405–426.

(8) Li, X. (2011) Infiltration of Nicotiana benthamiana Protocol for Transient Expression via Agrobacterium. BIO-Protoc. 1.

(9) Sauret-Güeto, S., Frangedakis, E., Silvestri, L., Rebmann, M., Tomaselli, M., Markel, K., Delmans, M., West, A., Patron, N. J., and Haseloff, J. (2020) Systematic Tools for Reprogramming Plant Gene Expression in a Simple Model, Marchantia polymorpha. ACS Synth. Biol. 9, 864–882.

(10) Annese, D., Romani, F., Grandellis, C., Ives, L., Frangedakis, E., Buson, F. X., Molloy, J. C., and Haseloff, J. (2025) Semi-automated workflow for high-throughput *Agrobacterium* - mediated plant transformation. Plant J. 122, e70118.

(11) Ishizaki, K., Nishihama, R., Ueda, M., Inoue, K., Ishida, S., Nishimura, Y., Shikanai, T., and Kohchi, T. (2015) Development of Gateway Binary Vector Series with Four Different Selection Markers for the Liverwort Marchantia polymorpha. PLOS ONE 10, e0138876.

(12) Tse, S. W., Annese, D., Romani, F., Guzman-Chavez, F., Bonter, I., Forestier, E., Frangedakis, E., and Haseloff, J. (2024) Optimising promoters and subcellular localisation for constitutive transgene expression in Marchantia polymorpha. Plant Cell Physiol. pcae063.

(13) Frangedakis, E., Guzman-Chavez, F., Rebmann, M., Markel, K., Yu, Y., Perraki, A., Tse, S. W., Liu, Y., Rever, J., Sauret-Gueto, S., Goffinet, B., Schneider, H., and Haseloff, J. (2021) Construction of DNA tools for hyper-expression in Marchantia chloroplasts. ACS Synth. Biol.

(14) Muyldermans, S. (2013) Nanobodies: Natural Single-Domain Antibodies. Annu. Rev. Biochem. 82, 775–797.

(15) Bannas, P., Hambach, J., and Koch-Nolte, F. (2017) Nanobodies and Nanobody-Based Human Heavy Chain Antibodies As Antitumor Therapeutics. Front. Immunol. 8, 1603.

(16) Harmsen, M. M., and De Haard, H. J. (2007) Properties, production, and applications of camelid single-domain antibody fragments. Appl. Microbiol. Biotechnol. 77, 13–22.

(17) Coherent Market Insights. (2023, September) Nanobodies Market Size, Trends and Forecast to 2030. Nanobodies Mark. Anal.

(18) Modarresi, M., Javaran, M. J., Shams-Bakhsh, M., Zeinali, S., Behdani, M., and Mirzaee, M. (2018) Transient expression of anti-VEFGR2 nanobody in Nicotiana tabacum and N. benthamiana. 3 Biotech 8, 484.

(19) Teh, Y.-H. A., and Kavanagh, T. A. (2010) High-level expression of Camelid nanobodies in Nicotiana benthamiana. Transgenic Res. 19, 575–586.

(20) De Meyer, T., Arcalis, E., Melnik, S., Maleux, K., Nolf, J., Altmann, F., Depicker, A., and Stöger, E. (2020) Seed-produced anti-globulin VHH-Fc antibodies retrieve globulin precursors in the insoluble fraction and modulate the Arabidopsis thaliana seed subcellular morphology. Plant Mol. Biol. 103, 597–608.

(21) De Wilde, K., De Buck, S., Vanneste, K., and Depicker, A. (2013) Recombinant Antibody Production in Arabidopsis Seeds Triggers an Unfolded Protein Response. Plant Physiol. 161, 1021–1033.

(22) Jobling, S. A., Jarman, C., Teh, M.-M., Holmberg, N., Blake, C., and Verhoeyen, M. E. (2003) Immunomodulation of enzyme function in plants by single-domain antibody fragments. Nat. Biotechnol. 21, 77–80.

(23) Fridy, P. C., Li, Y., Keegan, S., Thompson, M. K., Nudelman, I., Scheid, J. F., Oeffinger, M., Nussenzweig, M. C., Fenyö, D., Chait, B. T., and Rout, M. P. (2014) A robust pipeline for rapid production of versatile nanobody repertoires. Nat. Methods 11, 1253–1260.

(24) Wang, Z., Li, L., Hu, R., Zhong, P., Zhang, Y., Cheng, S., Jiang, H., Liu, R., and Ding, Y. (2021) Structural insights into the binding of nanobodies LaM2 and LaM4 to the red fluorescent protein mCherry. Protein Sci. Publ. Protein Soc. 30, 2298–2309.

(25) Yamagata, M., and Sanes, J. R. (2018) Reporter–nanobody fusions (RANbodies) as versatile, small, sensitive immunohistochemical reagents. Proc. Natl. Acad. Sci. 115, 2126–2131.

(26) Shaner, N. C., Campbell, R. E., Steinbach, P. A., Giepmans, B. N. G., Palmer, A. E., and Tsien, R. Y. (2004) Improved monomeric red, orange and yellow fluorescent proteins derived from Discosoma sp. red fluorescent protein. Nat. Biotechnol. 22, 1567–1572.

(27) Schriek, A. I., van Haaren, M. M., Poniman, M., Dekkers, G., Bentlage, A. E. H., Grobben, M., Vidarsson, G., Sanders, R. W., Verrips, T., Geijtenbeek, T. B. H., Heukers, R., Kootstra, N. A., de Taeye, S. W., and van Gils, M. J. (2022) Anti-HIV-1 Nanobody-IgG1 Constructs With Improved Neutralization Potency and the Ability to Mediate Fc Effector Functions. Front. Immunol. 13.

(28) Jumper, J., Evans, R., Pritzel, A., Green, T., Figurnov, M., Ronneberger, O., Tunyasuvunakool, K., Bates, R., Žídek, A., Potapenko, A., Bridgland, A., Meyer, C., Kohl, S. A. A., Ballard, A. J., Cowie, A., Romera-Paredes, B., Nikolov, S., Jain, R., Adler, J., Back, T., Petersen, S., Reiman, D., Clancy, E., Zielinski, M., Steinegger, M., Pacholska, M., Berghammer, T., Bodenstein, S., Silver, D., Vinyals, O., Senior, A. W., Kavukcuoglu, K., Kohli, P., and Hassabis, D. (2021) Highly accurate protein structure prediction with AlphaFold. Nature 596, 583–589.

(29) Damaj, M. B., Jifon, J. L., Woodard, S. L., Vargas-Bautista, C., Barros, G. O. F., Molina, J., White, S. G., Damaj, B. B., Nikolov, Z. L., and Mandadi, K. K. (2020) Unprecedented enhancement of recombinant protein production in sugarcane culms using a combinatorial promoter stacking system. Sci. Rep. 10, 13713.

(30) Hopp, T. P., Prickett, K. S., Price, V. L., Libby, R. T., March, C. J., Pat Cerretti, D., Urdal, D. L., and Conlon, P. J. (1988) A Short Polypeptide Marker Sequence Useful for Recombinant Protein Identification and Purification. Bio/Technology 6, 1204–1210.

(31) Coyle, B. L., and Baneyx, F. (2014) A cleavable silica-binding affinity tag for rapid and inexpensive protein purification. Biotechnol. Bioeng. 111, 2019–2026.

(32) Suzuki, H., Kato, H., Iwano, M., Nishihama, R., and Kohchi, T. (2023) Auxin signaling is essential for organogenesis but not for cell survival in the liverwort Marchantia polymorpha. Plant Cell 35, 1058–1075.

(33) Ismaili, A., Jalali-Javaran, M., Rasaee, M. J., Rahbarizadeh, F., Forouzandeh-Moghadam, M., and Memari, H. R. (2007) Production and characterization of anti-(mucin MUC1) single-domain antibody in tobacco (Nicotiana tabacum cultivar Xanthi). Biotechnol. Appl. Biochem. 47, 11–19.

(34) Richard, G., Meyers, A. J., McLean, M. D., Arbabi-Ghahroudi, M., MacKenzie, R., and Hall, J. C. (2013) In vivo neutralization of α-cobratoxin with high-affinity llama single-domain antibodies (VHHs) and a VHH-Fc antibody. PloS One 8, e69495.

(35) Mardanova, E. S., Blokhina, E. A., Tsybalova, L. M., Peyret, H., Lomonossoff, G. P., and Ravin, N. V. (2017) Efficient Transient Expression of Recombinant Proteins in Plants by the Novel pEff Vector Based on the Genome of Potato Virus X. Front. Plant Sci. 8.

(36) Garabagi, F., Gilbert, E., Loos, A., McLean, M. D., and Hall, J. C. (2012) Utility of the P19 suppressor of gene-silencing protein for production of therapeutic antibodies in Nicotiana expression hosts. Plant Biotechnol. J. 10, 1118–1128.

(37) Matsuo, K., and Atsumi, G. (2019) CRISPR/Cas9-mediated knockout of the RDR6 gene in Nicotiana benthamiana for efficient transient expression of recombinant proteins. Planta 250, 463–473.

(38) Matsuo, K. (2022) CRISPR/Cas9-mediated knockout of the DCL2 and DCL4 genes in Nicotiana benthamiana and its productivity of recombinant proteins. Plant Cell Rep. 41, 307–317.

(39) Romani, F., Sauret-Güeto, S., Rebmann, M., Annese, D., Bonter, I., Tomaselli, M., Dierschke, T., Delmans, M., Frangedakis, E., Silvestri, L., Rever, J., Bowman, J. L., Romani, I., and Haseloff, J. (2024) The landscape of transcription factor promoter activity during vegetative development in Marchantia. Plant Cell koae053.

(40) Stark, J. C., Huang, A., Nguyen, P. Q., Dubner, R. S., Hsu, K. J., Ferrante, T. C., Anderson, M., Kanapskyte, A., Mucha, Q., Packett, J. S., Patel, P., Patel, R., Qaq, D., Zondor, T., Burke, J., Martinez, T., Miller-Berry, A., Puppala, A., Reichert, K., Schmid, M., Brand, L., Hill, L. R., Chellaswamy, J. F., Faheem, N., Fetherling, S., Gong, E., Gonzalzles, E. M., Granito, T., Koritsaris, J., Nguyen, B., Ottman, S., Palffy, C., Patel, A., Skweres, S., Slaton, A., Woods, T., Donghia, N., Pardee, K., Collins, J. J., and Jewett, M. C. (2018) BioBits^TM^ Bright: A fluorescent synthetic biology education kit. Sci. Adv. 4, eaat5107.

(41) Boehm, C. R., Ueda, M., Nishimura, Y., Shikanai, T., and Haseloff, J. (2016) A Cyan Fluorescent Reporter Expressed from the Chloroplast Genome of Marchantia polymorpha. Plant Cell Physiol. 57, 291–299.

(42) Dudley, Q. M., Cai, Y.-M., Kallam, K., Debreyne, H., Carrasco Lopez, J. A., and Patron, N. J. (2021) Biofoundry-assisted expression and characterization of plant proteins. Synth. Biol. 6, ysab029.

(43) Schulte, R., Talamas, J., Doucet, C., and Hetzer, M. W. (2008) Single Bead Affinity Detection (SINBAD) for the Analysis of Protein-Protein Interactions. PLOS ONE 3, e2061.

(44) Kim, J. M., Seong, B. L., and Lim, D.-K. (2020) Bead based facile assay for sensitive quantification of native state green fluorescent protein. RSC Adv. 10, 13095–13099.

