## Additional information on experimental details, plasmids and primers used for "*Marchantia polymorpha* as a simple platform for plant-based production of functional nanobodies"

### Production of an anti-mCherry Nanobody-Turquoise2 fusion in *Marchantia polymorpha*

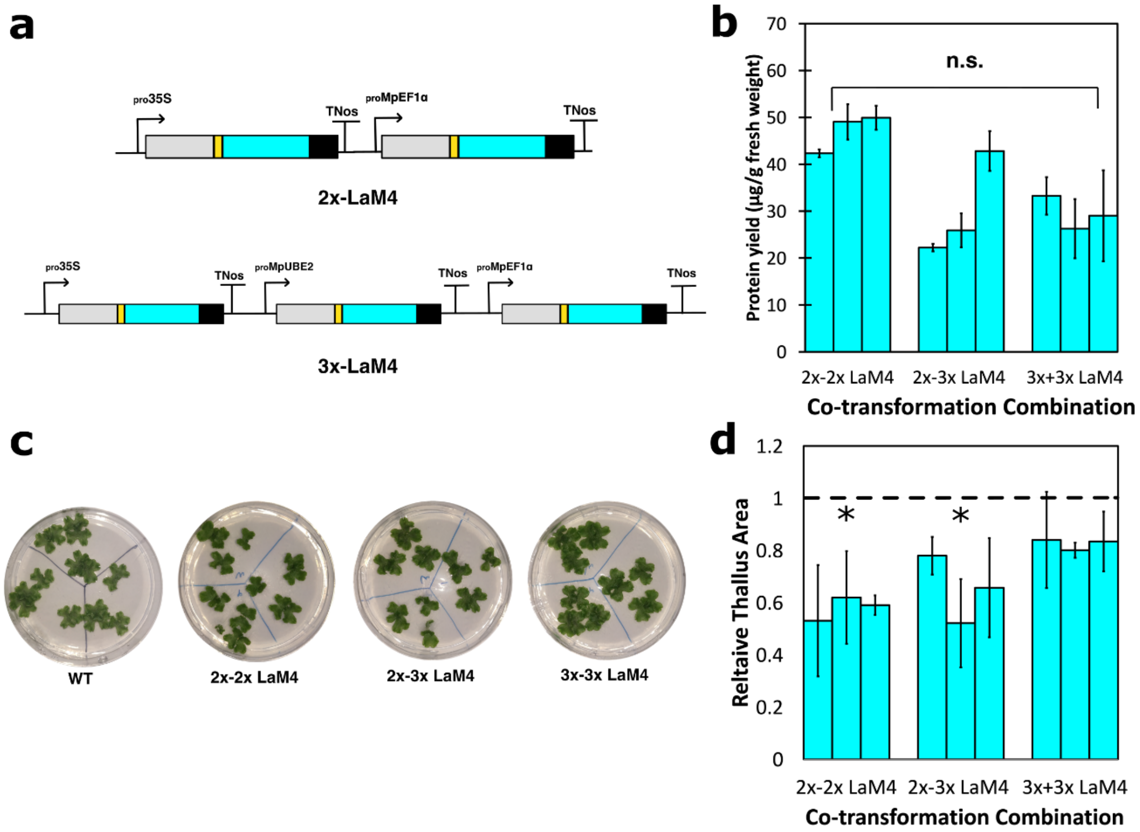


***Figure S1**: Expression level of LaM4-mTurquoise2-7xHis and relative size estimation of plants from different in different co-transformation lines. **a**: Schematic representation of different co-transformation lines constructs. **b**: LaM4-mTurquoise2-7xHis expressed in different co-transformation lines in 3-week-old plants, in µg/g fresh weight. **c**: Images of 2-week-old WT and different co-transformation lines plants grown in 9cm petri-dishes in 0.5× Gamborg B-5 basal medium. The plates were divided into three portions, and on each portion grew 3 gemmae derived from the same independent transformant of that promoter construct. **d**: Relative thallus size (the ratio of sizes of co-transformation lines plants: average size of WT plants; dash line indicates WT size) of different promoter line plants (2-week-old). In **b** and **d**, each bar represents an independent transformant for that promoter construct, and error bars represents standard deviation between 3 biological replicates. Asterisk above the bars indicate statistically significant differences between the different co-transformation lines(Dunn's Test; $p < 0.05$).*

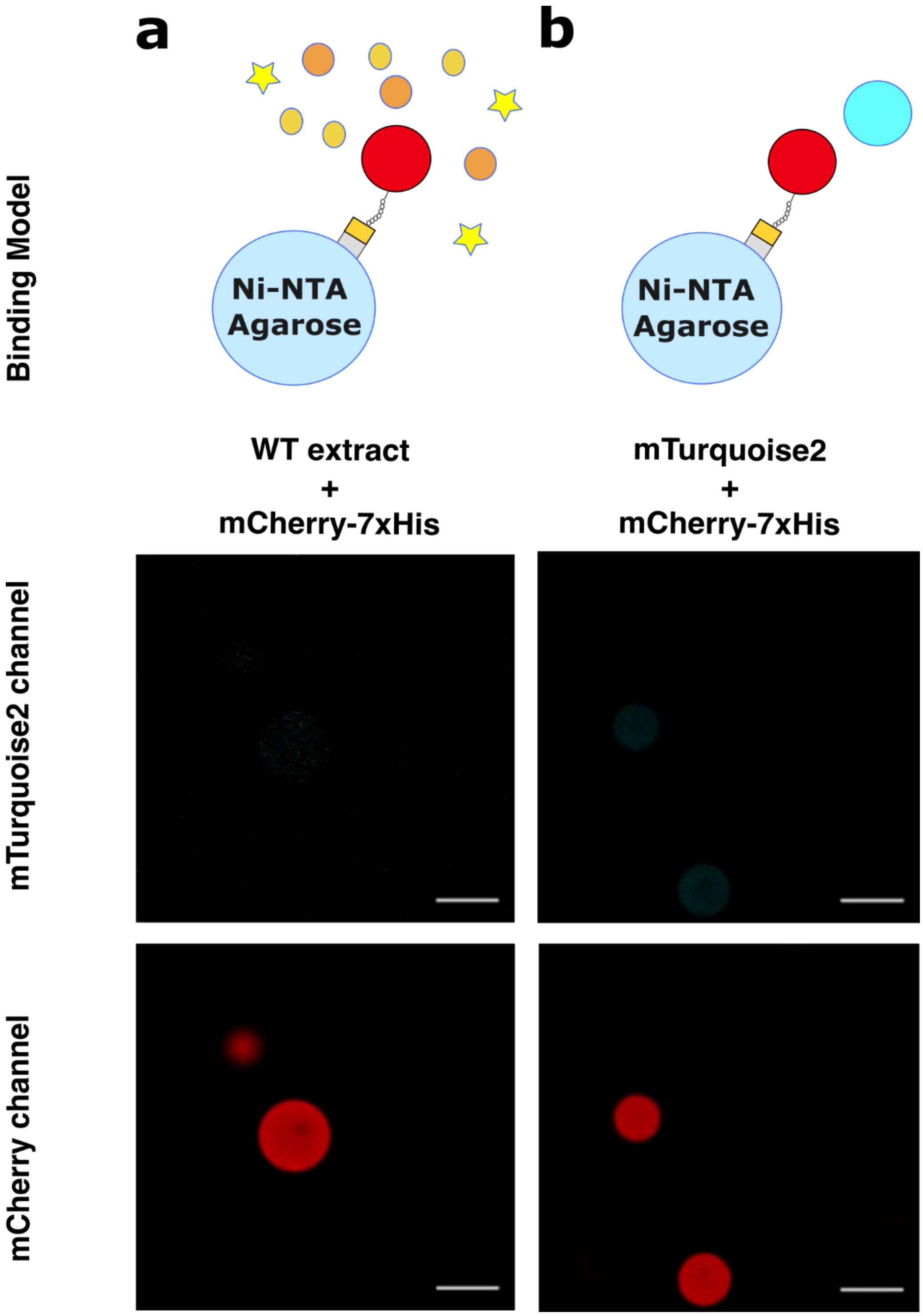


***Figure S2***: *Negative controls for Ni-NTA anti-His agarose beads with plant extracts. a: mCherry-7xHis incubated with WT Marchantia extract and Ni-NTA agarose beads. b: LaM4-mTurquoise2-3xFLAG incubated with mCherry-7xHis, mTurquoise2 and Ni-NTA agarose beads. mTurquoise2 signal shown as cyan and mCherry signal shown in red. Scale = 100µm.*

| Name | Plasmid details | Purpose | Source |
|---|---|---|---|
| pmTurquoise2 | $_{pro}$T7:mTurquoise2 | E coli expression of tagless mTurquoise2 | This paper |
| pCRB SREI $_{6his}$ | $_{pro}$T7:mTurquoise2-6xHis | E coli expression of mTurquoise2-6x-His | Boehm et al., (please refer to main text for citation) |
| pJL1-mCherry | $_{pro}$T7:mCherry | E coli expression of tagless mCherry | Addgene (please refer to main text for citation) |
| pmCherry-7xHis | $_{pro}$T7:mCherry-7xHis | E coli expression of mCherry-7xHis | This paper |
| pT7-Nb | $_{pro}$T7: LaM4-mTurquoise2-7xHis | E coli expression of LaM4-mTurquoise2-7xHis | This paper |
| $_{pro}$MpEF1$\alpha$: Nb-7xHis | $_{pro}$MpEF1$\alpha$: LaM4-mTurquoise2-7xHis | In planta expression of LaM4-mTurquoise2-7xHis | This paper |
| $_{pro}$MpUBE2: Nb-7xHis | $_{pro}$MpUBE2: LaM4-mTurquoise2-7xHis | | |
| $_{pro}$35S: Nb-7xHis | $_{pro}$35S: LaM4-mTurquoise2-7xHis | | |
| $_{pro}$35S×2:Nb-7xHis | $_{pro}$35S×2: LaM4-mTurquoise2-7xHis | | |
| 2x-Hyg-Nb-7xHis | HygR-$_{pro}$35S: LaM4-mTurquoise2-7xHis-spacer- $_{pro}$MpEF1$\alpha$: LaM4-mTurquoise2-7xHis | | |
| 2x-CsR-Nb-7xHis | CsR-$_{pro}$35S: LaM4-mTurquoise2-7xHis-spacer- $_{pro}$MpEF1$\alpha$: LaM4-mTurquoise2-7xHis | | |
| 2x-35Sx2-Nb-7xHis | HygR-$_{pro}$35S×2: LaM4-mTurquoise2-7xHis- spacer- $_{pro}$MpEF1$\alpha$: LaM4-mTurquoise2-7xHis | | |
| 3x-Hyg-Nb-7xHis | HygR-$_{pro}$35S: LaM4-mTurquoise2-7xHis-$_{pro}$MpUBE2: LaM4-mTurquoise2-7xHis-$_{pro}$MpEF1$\alpha$: LaM4-mTurquoise2-7xHis | | |

| Name | Plasmid details | Purpose | Source |
|---|---|---|---|
| 3x-CsR-Nb-7xHis | CsR-$_{pro}$35S: LaM4-mTurquoise2-7xHis--<br>$_{pro}$MpUBE2: LaM4-mTurquoise2-7xHis-<br>$_{pro}$MpEF1$\alpha$: LaM4-mTurquoise2-7xHis | In planta expression of LaM4-mTurquoise2-7xHis/alternative tags | This paper |
| $_{pro}$35S×2:Nb-3xFLAG | $_{pro}$35S×2: LaM4-mTurquoise2-3xFLAG | | |
| $_{pro}$35S×2:Nb-Car9 | $_{pro}$35S×2: LaM4-mTurquoise2-Car9 | | |
| $_{pro}$35S×2:Nb-cCar9 | $_{pro}$35S×2: LaM4-mTurquoise2-cCar9 | | |

***Table S1****: All plasmids used in the study.*

| Name | Primer sequence 5'->3' | Purpose |
| --- | --- | --- |
| mTurquoise2-F | TTTGGTCTCTAATGGTGAGCAAGGGCGAGGAGCTGTT | Amplification of mCherry-7xHis fragment for pmTurquoise2 |
| mTurquoise2-R | AAAGGTCTCTAAGCTTACTTGTACAGCTCGTCCATGCCGAGA | Amplification of mCherry-7xHis fragment for pmTurquoise2 |
| mTurquoise2-7xHis-F | TTGGTCTCTGTGAGCAAGGGCGAGGAGCTGTT | Amplification of the mTurquoise2-7xHis fragment for all LaM4-mTurquoise2-7xHis plasmids and the mTurquoise fragment for all LaM4-mTurquoise2 plasmids with alternative tags |
| mTurquoise2-7xHis-R | AAGGTCTCTAAGCTTAATGATGGTGATGGTGATGGTGACCAGAAGACTTGTACAGCTCGTCCATGCCGAGA | Amplification of the mTurquoise2-7xHis fragment for all LaM4-mTurquoise2-7xHis plasmids |
| mTurquoise2-CDSI-R | AAGGTCTCTCGAAGCCTTGTACAGCTCGTCCATGCCGAGA | Amplification of the mTurquoise fragment for all LaM4-mTurquoise2 plasmids with alternative tags |
| mCherry-7xHis-F | TTGGTCTCGAATGGTGAGCAAGGGCGAGGAGG | Amplification of mCherry-7xHis fragment for pmCherry-7xHis |
| mCherry-7xHis-R | TTTGGTCTCGAAGCTTAATGATGGTGATGGTGATGGTGACCAGAAGAAGCCTTGTACAGCTCGTCCATGCC | Amplification of mCherry-7xHis fragment for pmCherry-7xHis |
| 3xFLAG-F | TTCGAACAGCGACTACAAAGACCATGACGGTGATTATAAAGATCATGACATCGACTACAAGGATGACGATGACAAGTAA | Creating 3xFLAG part as a primer dimer |

| | | |
|---|---|---|
| 3xFLAG-R | AAGCTTACTTGTCATCGTCATCCTTGTAGTCGATGTCATGATCTTTATAATCACCGTCATGGTCTTTGTAGTCGCTGTT | Creating 3xFLAG part as a primer dimer |
| Car9-F | TTCGGGTGGTGGGAGTGATAGCGCGCGGCTTTAAAAAACCGGGAAAGAGGTAA | Creating Car9 part as a primer dimer |
| Car9-R | AGCTTACCTCTTTCCCGGTTTTTTAAAGCCGCGCGCTATCACTCCCACCACC | Creating Car9 part as a primer dimer |
| cCar9-F | TTCGAAGAAAGGTGGTGGGAGTGATAGCGCGCGGCTTTAAAAAACCGGGAAAGAGGTAA | Creating E. coli OmpT cleavable Car9 part as a primer dimer |
| cCar9-R | AAGCTTACCTCTTTCCCGGTTTTTTAAAGCCGCGCGCTATCACTCCCACCACCTTTCTT | Creating E. coli OmpT cleavable Car9 part as a primer dimer |

***Table S2:*** *All primers used in the study.*
